# PRDM1 controls the sequential activation of neural, neural crest and sensory progenitor determinants by regulating histone modification

**DOI:** 10.1101/607739

**Authors:** Ravindra S. Prajapati, Mark Hintze, Andrea Streit

## Abstract

During early embryogenesis, the ectoderm is rapidly subdivided into neural, neural crest and sensory progenitors. How the onset of lineage-specific determinants and the loss of pluripotency markers are temporally and spatially coordinated in vivo remains an open question. Here we identify a critical role for the transcription factor PRDM1 in the orderly transition from epiblast to defined neural lineages. Like pluripotency factors, *PRDM1* is expressed in all epiblast cells prior to gastrulation, but lost as they begin to differentiate. We show that, unlike pluripotency factors, PRDM1 is initially required for the activation of neural, neural crest and sensory progenitor specifiers and for the downregulation of pluripotency-associated genes. In vivo chromatin immunoprecipitation reveals stage-specific binding of PRDM1 to regulatory regions of neural and sensory progenitor genes, PRDM1-dependent recruitment of the histone demethylase Kdm4a to these regions and associated removal of repressive histone marks. Once lineage determinants become expressed, they repress *PRDM1*, and our data suggest that *PRDM1* downregulation is required for cells to maintain their identity. Thus, PRDM1 mediates chromatin modifications that directly control neural and sensory progenitor genes, and its activities switch from an activator at early stages to a repressor once neural fates have been established.

## INTRODUCTION

In human and mouse embryonic stem cells, exit from pluripotency and entry into differentiation programmes is accompanied by dramatic changes in the chromatin landscape (Andrey and Mundlos, 2017; Habibi and Stunnenberg, 2017; Kalkan and Smith, 2014; Kim et al., 2008; Li and Izpisua Belmonte, 2018; Schlesinger and Meshorer, 2019; Surani et al., 2007; Theunissen and Jaenisch, 2017). While cells gradually lose the expression of pluripotency genes, developmental genes are primed for activation by changes in histone tail modifications. Subsequently, cross-repressive interactions between different transcription factors are thought to establish mutually exclusive fates. However, recent evidence suggests that pluripotency and differentiation networks overlap to varying degrees prior to final fate specification. A major challenge remaining is how to translate these processes defined in vitro to the developing embryo, where exit form pluripotency is not only controlled in time but is also synchronised with cell and tissue rearrangements that lay down the body plan (Habibi and Stunnenberg, 2017; Posfai et al., 2014; Rossant and Tam, 2017; Theunissen and Jaenisch, 2017; Wamaitha and Niakan, 2018).

Amniote epiblast cells have the potential to form all embryonic lineages and a small network of transcription factors including *PouV/Oct4, Nanog* and *Sox2* or *-3* (Dunn et al., 2014; Kalkan and Smith, 2014; Kim et al., 2008; Rossant and Tam, 2017), and *ERNI/Ens1* in birds (Fernandez-Tresguerres et al., 2010; Jean et al., 2015; Trevers et al., 2017) maintains them in a pluripotent undifferentiated state. Like in humans, the chick epiblast is a flat disc and this morphology is ideal to visualise rapid changes in gene expression in time and space as epiblast cells activate lineage-specific programmes. During gastrulation the epiblast is transformed into three germ layers with non-ingressing epiblast cells forming the ectoderm, which generates precursors for the central and peripheral nervous system in quick succession starting from the epiblast centre and progressing towards its periphery (Basch et al., 2006; Litsiou et al., 2005; Puelles et al., 2005; Streit et al., 1998; Streit and Stern, 1999; Stuhlmiller and García-Castro, 2012; Trevers et al., 2017; for review: Pla and Monsoro-Burq, 2018; Streit, 2018). The definitive neural plate, the primordium for the central nervous system, forms centrally surrounding the organiser (Fernandez-Garre et al., 2002; Rex et al., 1997; Sanchez-Arrones et al., 2012; Streit et al., 1997; Uchikawa et al., 2003), while neural crest and sensory progenitor fates emerge slightly later from cells at the neural plate border (Basch et al., 2006; Ezin et al., 2009; Khudyakov and Bronner-Fraser, 2009; Litsiou et al., 2005; for review: Pla and Monsoro-Burq, 2018; Simoes-Costa and Bronner, 2013, 2015; Streit, 2018). Neural plate border cells contain progenitors for neural, neural crest and sensory placode lineages and uniquely retain much of the pluripotency network throughout gastrula and neural plate stages endowing them with stem cell like properties (Buitrago-Delgado et al., 2015; Buitrago-Delgado et al., 2018; Hintze et al., 2017; Trevers et al., 2017). As different fates are allocated, cells lose the expression of pluripotency markers, while activating fate specifiers like the definitive neural marker *Sox2*, the neural crest marker *Foxd3* and the sensory progenitor genes *Six1* and *Eya1/2* (Buitrago-Delgado et al., 2015; Buitrago-Delgado et al., 2018; Hintze et al., 2017; Trevers et al., 2017). How is the sequential transition towards lineage determination controlled?

While some of the signalling events have been identified, we know relatively little about the cell intrinsic mechanisms that coordinate the temporal and spatial order in which neural, neural crest and sensory progenitors are specified. In chick, the coiled-coil domain proteins ERNI and BERT play an important role in controlling the timing of *Sox2* expression in the neural plate through its N2 enhancer (Papanayotou et al., 2008). At early gastrulation stages, the N2 enhancer is occupied by the chromatin remodelling enzyme Brm and the nuclear factors Geminin and ERNI, which in turn recruits transcriptional repressors to prevent premature activation. Towards the end of gastrulation, *BERT* expression is initiated and replaces ERNI in this complex thus allowing *Sox2* to be expressed. Identification of the gene networks that regulate neural crest and sensory progenitor specification reveals that both fates are initially under the control of neural plate border genes, which act in different combinations to confer neural crest or sensory progenitor identity. Foxd3 is a key neural crest determination factor (Kos et al., 2001; Lukoseviciute et al., 2018; Mundell and Labosky, 2011; Sasai et al., 2001; Simoes-Costa et al., 2012; Teng et al., 2008), and its enhancers are regulated by a combination of Pax3/7, Msx1 and Zic1 as well as the pluripotency factors Sox2, Nanog and Oct3/4 (Fujita et al., 2016; Simoes-Costa et al., 2012). In contrast, the sensory progenitor determinant Six1 is directly controlled by Dlx5/6, negatively regulated by Msx1, and probably indirectly by Gata3 and Tfap2a (Kwon et al., 2010; Pieper et al., 2012; Sato et al., 2010). However, how cells transit from an early epiblast state associated with pluripotency to activating cell fate specific programmes in a temporal and spatial order remains unclear.

Here we identify the transcription factor PRDM1 as a key component for the orderly transition from a pluripotency-like state to defined neural lineages. At its N-terminus PRDM1 contains a methyltransferase-like PR/SET domain, which lacks enzymatic activity, while five C-terminal C2H2 zinc fingers mediate DNA binding and recruitment of chromatin modifying enzymes (for review: Bikoff et al., 2009). Mostly acting as a transcriptional repressor (Ancelin et al., 2006; Gyory et al., 2004; Kurimoto et al., 2015; Ren et al., 1999), it is known for its critical role in B-and T-lymphocyte differentiation, germ cell fate determination, as well as in limb, heart and pharyngeal development (Kallies and Nutt, 2007; Magnusdottir et al., 2013; Nutt et al., 2007; Ohinata et al., 2005; Robertson et al., 2007; Saitou et al., 2005; Senft et al., 2019; Shaffer et al., 2002; Vincent et al., 2005). In zebrafish, PRDM1 also controls neural crest cell formation by directly regulating *Foxd3* (Hernandez-Lagunas et al., 2005; Olesnicky et al., 2010; Powell et al., 2013). Here we show that in chick *PRDM1* expression is remarkably similar to that of the pluripotency associated gene *ERNI* (Streit et al., 2000): both are highly expressed in the pre-gastrula epiblast together with other pluripotency associated transcripts, but are gradually lost form ectodermal cells as neural, neural crest and sensory progenitor lineages are established. We find that *PRDM1* is initially required for the loss of pluripotency markers and for the acquisition of all neural, neural crest and sensory progenitor identity. PRDM1 acts as a transcriptional activator by recruiting Kdm4a to the promoter regions of neural and sensory progenitor genes, which in turn removes repressive histone marks to facilitate their expression in a stage-specific manner. Subsequently, *PRDM1* downregulation is mediated by specifiers of each neural fate - Sox2, Foxd3 and Six1. This is required to maintain their expression since PRDM1 function has now switched to repress neural, neural crest and sensory progenitor identity. Therefore, during early ectoderm development PRDM1 has three distinct activities: it represses pluripotency associated genes, directly activates neural determinants, and must later be lost to allow progression towards definitive neural, neural crest and sensory progenitor identity.

## RESULTS

### Loss of *PRDM1* expression accompanies neural, neural crest and sensory progenitor specification

We have recently found that the transcription factor *PRDM1* is highly enriched in the pre-streak epiblast together with other pluripotency markers like *ERNI, Sox3* and *Nanog*. PRDM1 is an important regulator of cell fate decisions (Kallies and Nutt, 2007; Magnusdottir et al., 2013; Nutt et al., 2007; Ohinata et al., 2005; Robertson et al., 2007; Saitou et al., 2005; Senft et al., 2019; Shaffer et al., 2002; Vincent et al., 2005), and we therefore compared its expression during early chick development with those of pluripotency, neural, neural crest and sensory progenitor makers. *PRDM1* expression begins prior to primitive streak formation, but after the onset of *ERNI, Sox3* and *Nanog* (Jean et al., 2015; Streit et al., 2000; Trevers et al., 2017) around stage EG XIII (Fig. 1A, B, M, N, a, b, m, n); all four transcripts are expressed almost identically in the entire epiblast. As the neural plate marker *Sox2* is activated (Fig. 1 G, H, g, h) *PRDM1* expression decreases in Sox2^+^ cells and becomes gradually confined to the sensory progenitor domain at HH5 (Fig. 1C, D, c, d). For a short time, *PRDM1* is co-expressed with the sensory progenitor markers *Six1* and *Eya2* (Fig. 1 K, L, k, l), but is gradually lost as sensory placodes are specified (Supplemental Fig. 1). These dynamic changes of *PRDM1* expression are reminiscent of that of *ERN1* and other pluripotency markers (Jean et al., 2015; Lavial et al., 2007; Streit et al., 2000; Trevers et al., 2017), which like *PRDM1* are downregulated as cells are specified as neural and sensory progenitors.

**Figure 1.**
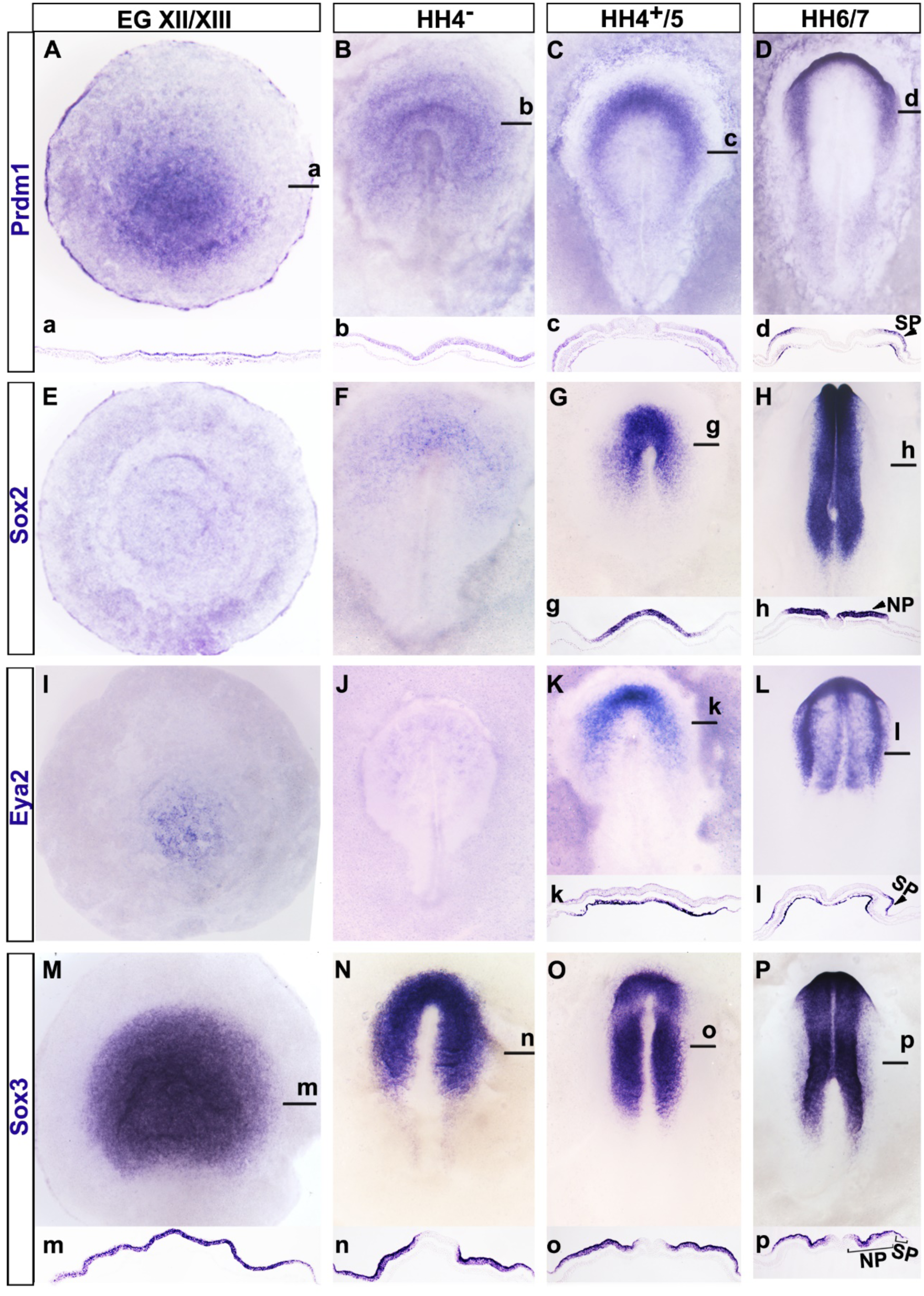
Expression of *PRDM1, Sox2, Eya2* and *Sox3* in the early chick embryo. A-D. *PRDM1* is broadly expressed in the epiblast at pre-primitive streak stages (A, a) and at primitive streak stages (B, b), but is downregulated as the neural plate is specified (C, c). At headfold stages (D, d) *PRDM1* is confined to sensory progenitors. E-H. *Sox2* is not expressed at pre-streak stages (E) and starts to be expressed in the ectoderm surrounding the organiser at primitive streak stages (F). As the neural plate forms (G, g) its expression increases and it is confined to the neural plate at headfold stages (H, h; NP). I-L. *Eya2* is expressed in the hypoblast before primitive streak formation (I), and not expressed at primitive streak stages (J). At late primitive streak stages *Eya2* is expressed in the mesoderm, but absent from the ectoderm (K, k). At headfold stages (L, l) it continues to be expressed in the head mesoderm and in the ectoderm is expressed in sensory progenitors (SP). M. – P. *Sox3* is broadly expressed in the pre-streak (M, m) and primitive streak stage epiblast (N, n). At head process (O, o) and headfold (P, p) stages it is strongly expressed in the neural plate (NP) and weaker in sensory progenitors (SP). a-d, g, h, k, l m-p show sections through the same embryos as shown in A-D, G, H, K, L, M-P at the level of the black lines.

### PRDM1 is required for neural, neural crest and sensory progenitor fates and for the loss of pluripotency markers at primitive streak stages

Acting in a protein complex, PRDM1 recruits histone modifying enzymes to target genes and is generally associated with transcriptional repression (Ancelin et al., 2006; Gyory et al., 2004; Kurimoto et al., 2015; Ren et al., 1999; for review: Bikoff et al., 2009; Mzoughi et al., 2016). Its expression akin to that of pluripotency markers suggests that it may maintain cells in a pluripotent state and prevent lineage specification. To test this hypothesis, we used a loss of function approach and assessed changes in gene expression initially focusing on sensory progenitors. We knocked down *PRDM1* using two antisense oligonucleotides (aONs) targeting intron-exon junctions (Supplemental Fig. S2A). Two different aONs (alone or together) or control ONs were electroporated broadly into the epiblast of early primitive streak stage chick embryos (HH3) and targeted sensory progenitors were harvested at HH6. As a first step to assess the effect of PRDM1 knockdown on many downstream targets, we used multiplex NanoString nCounter to examine transcript levels of 382 genes including markers for pre-streak epiblast and different ectodermal fates (Supplemental Table1 and Supplemental Fig. 2B). Surprisingly, we find that the sensory progenitor markers *Eya2, Six1* and *Irx1*, as well as their upstream regulators *Gata3, Dlx5* and -*6* are significantly downregulated, as is the neural crest marker *Snail2*. In contrast, the pluripotency associated transcripts *ERNI, Nanog* and *Eomes* are upregulated, as are other genes expressed broadly in the pre-streak epiblast like *MafA* (Supplemental Figure 2B). These findings suggest that PRDM1 may be required for the loss of pluripotency markers and for the initiation of lineage specification.

To corroborate our NanoString results and to gain spatial information on changes in gene expression not only in sensory progenitors, but also in neural and neural crest cells we performed in situ hybridisation after PRDM1 knockdown. Embryos were again electroporated with control or experimental ONs at early streak stages (HH3) prior to the onset of definitive neural, neural crest and sensory progenitor markers. After 16–24hrs, we assessed *Sox2* (neural), *Foxd3* (neural crest) and *Six1* and *Eya2* (sensory progenitor). We find that *PRDM1* knockdown diminishes the expression of all four transcripts in aON targeted cells (*Sox2*: n=8/10; *Foxd3*: n=4/6; *Six1*: n=7/10; *Eya2*: n=8/10; Fig. 2F-I, f-i), while control ONs have no effect (Fig. 2A-D, a-d; Sox2: 0/8; Foxd3: 0/5; Six1: 0/5, Eya2: 0/9). PRDM1 is also required for the maintenance of *Dlx6* and *Gata3* expression, which are already expressed in the non-neural ectoderm at early primitive streak stages (Supplemental Fig. 2B-F, c-f). In contrast, *PRDM1* knock-down causes expansion or upregulation of the pluripotency markers *Sox3* and *ERNI* (Sox3 aON: n = 5/5; control: n = 0/5; Fig. 2 E, e, J, j; ERNI: Supplemental Fig. S2B). To control for the specificity of PRDM1 knockdown, we performed rescue experiment by co-electroporating PRDM1 aONs with a vector encoding full length PRDM1 and Cherry. This restores normal expression of *Sox2, Eya2* and *Sox3* (Supplemental Fig. S3 A-F). Together, these results suggest an unexpected role for PRDM1 at early primitive streak stages. While PRDM1 seems to repress stem cell-like properties, it is required for the activation of the neural, neural crest and sensory progenitor programmes.

**Figure 2.**
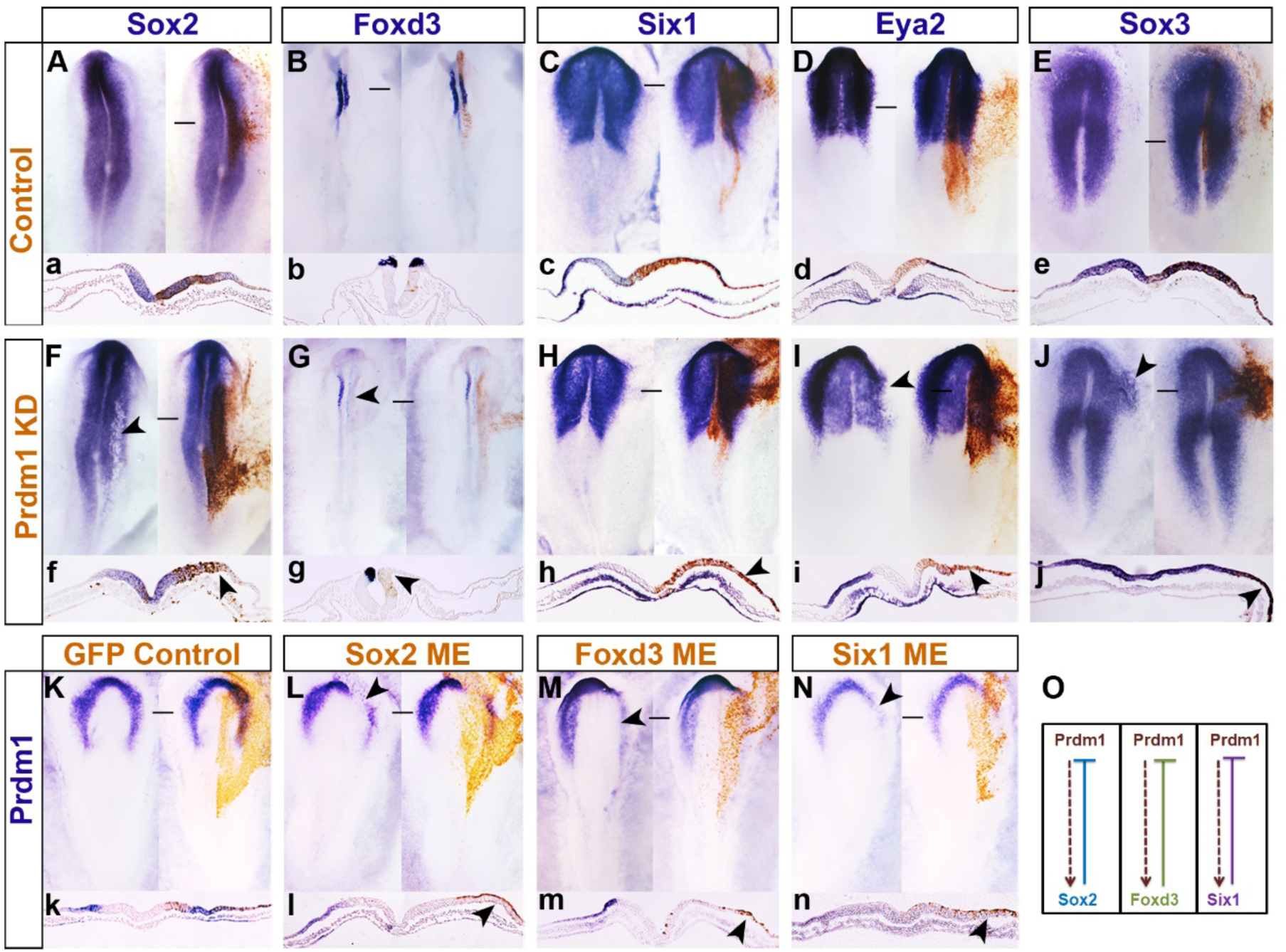
PRDM1 is required for neural, neural crest and sensory progenitor fates, but later repressed by fate determinants. Control (**A-E**, a-e) or PRDM1 targeting aOS (**F-J**, f-j) were electroporated into the epiblast of chick embryos at early primitive streak stages. At headfold stages the expression of *Sox2* (A, a, F, f), *Foxd3* (B, b, G, g), *Six1* (C, c, H, h), *Eya2* (D, d, I, i) and *Sox3* (E, e, J, j) was assessed by in situ hybridisation (blue). Fluorescein labelled OS are visualised by antibody staining in brown; panels on the left show each embryo before immunolabelling. a-j show sections through the same embryos as shown in A-J at the level of the black lines. Arrowheads in F-J and f-j point to changes in gene expression after PRDM1 knock down. **K**, k. Misexpression of GFP containing vectors (brown) do not affect the expression of *PRDM1* (blue); k shows section at the level of the black line in K. Panel on the left: embryo prior to immunolabelling for GFP (brown). Misexpression of Sox2 (**L**, l), Foxd3 (**M**, m) and Six1 (**N**, n) leads to downregulation of *PRDM1* (blue; arrowheads) in targeted cells (brown). l-n show sections of the same embryos shown in L-N at the level of the black line. O. PRDM1 is required for the expression of *Sox2, Foxd3* and *Six1* in the early epiblast, but is later repressed by these factors.

### Mutual repression between PRDM1 and Sox2, Foxd3 and Six1 maintains cell identity

While our results propose that *PRDM1* is necessary for specification of the entire nervous system, *PRDM1* expression is downregulated as soon as cell fate specifiers are expressed. Is PRDM1 loss required for cells to maintain lineage specification? To test this, we electroporated full length PRDM1 at primitive streak stages (HH4^−^; this leads to protein production about 3-5 hrs later at HH4/5). We find that at HH6/7 *Sox2* (5/5) and *Eya2* (4/5) are lost in *PRDM1* expressing cells (Supplemental Fig. 4B, D, b, d) suggesting that its downregulation is required for cells to maintain definitive neural and sensory progenitor identity. What mechanism is responsible for the downregulation *PRDM1*? In zebrafish, *PRDM1* is necessary for neural crest cell formation and it regulates *Foxd3* directly. However, once it starts to be expressed, Foxd3 represses *PRDM1* (Powell et al., 2013). We therefore tested whether a similar regulatory relationship exists in chick. HH3^+^ chick embryos were electroporated with Sox2, Foxd3 and Six1 full length expression constructs and the expression of PRDM1 was assessed at HH6/7 by in situ hybridisation. Misexpression of either Sox2 (9/9), Foxd3 (13/13) or Six1 (7/8) in the ectoderm leads to loss of *PRDM1*, while GFP controls are normal (Fig. 2K-N, k-n). Therefore, once ectodermal cells begin to acquire their unique identity at late primitive streak stages, repression of *PRDM1* by Sox2, Foxd3 and Six1 allows lineage progression towards neural, neural crest and sensory progenitors. These data suggest that PRDM1 has at least two distinct activity phases, first prior to the activation of the neural programmes and second during further specification.

### PRDM1 occupies Sox2 and Eya2 promoter regions at distinct stages

Our results show that PRMD1 is required early for the expression of general neural progenitor genes. Does it act by direct interaction with regulatory or promoter regions? We find PRDM1 motifs within 2kb upstream of the transcription start site (TSS) of Sox2, Foxd3, Six1 and Eya2, and of neural plate border genes like Dlx5/6, Msx2 and TFAP2a/e and transcripts expressed in a subset of sensory progenitors (e.g. Pax6, SSTR5, Dmbx1) (Supplemental Table 2). To assess whether PRDM1 occupies these sites in vivo, we focused on Sox2 and Eya2 as key factors required for neural plate and sensory progenitor specification and performed PRDM1 chromatin immunoprecipitation (ChIP) followed by qPCR (Fig. 3A-D). For this analysis, we dissected four distinct embryonic territories representing different cell states (Fig. 3B): epiblast cells prior to primitive streak formation (cEpi: *PRDM1*^*+*^, *Sox2*^*-*^, *Eya2*^*-*^), the neural plate border (NPB: *PRDM1*^*+*^, *Sox2*^*-*^, *Eya2*^*-*^) and early neural plate (eNP: *PRDM1*^*+*^, *Sox2*^*+*^, *Eya2*^*-*^) from HH4^−^, and the sensory progenitor domain (SP: *PRDM1*^*+*^, *Sox2*^*-*^, *Eya2*^*+*^) from HH6. ChIP-qPCR reveals a significant enrichment of PRDM1 upstream of the Sox2 and Eya2 TSS when compared to IgG controls, but only in early neural plate and sensory progenitor cells, respectively (Fig. 3C, D). As a negative control we examined the TSS of the histone demethylase Kdm4a, which is ubiquitously expressed in epiblast cells at all the stages tested (Strobl-Mazzulla et al., 2010) and lacks a PRDM1 motif. We do not observe PRDM1 binding (Fig. 3I). Thus, when *Sox2* and *Eya2* begin to be expressed PRDM1 is bound close to their TSS suggesting that it may regulate their transcription directly.

**Figure 3.**
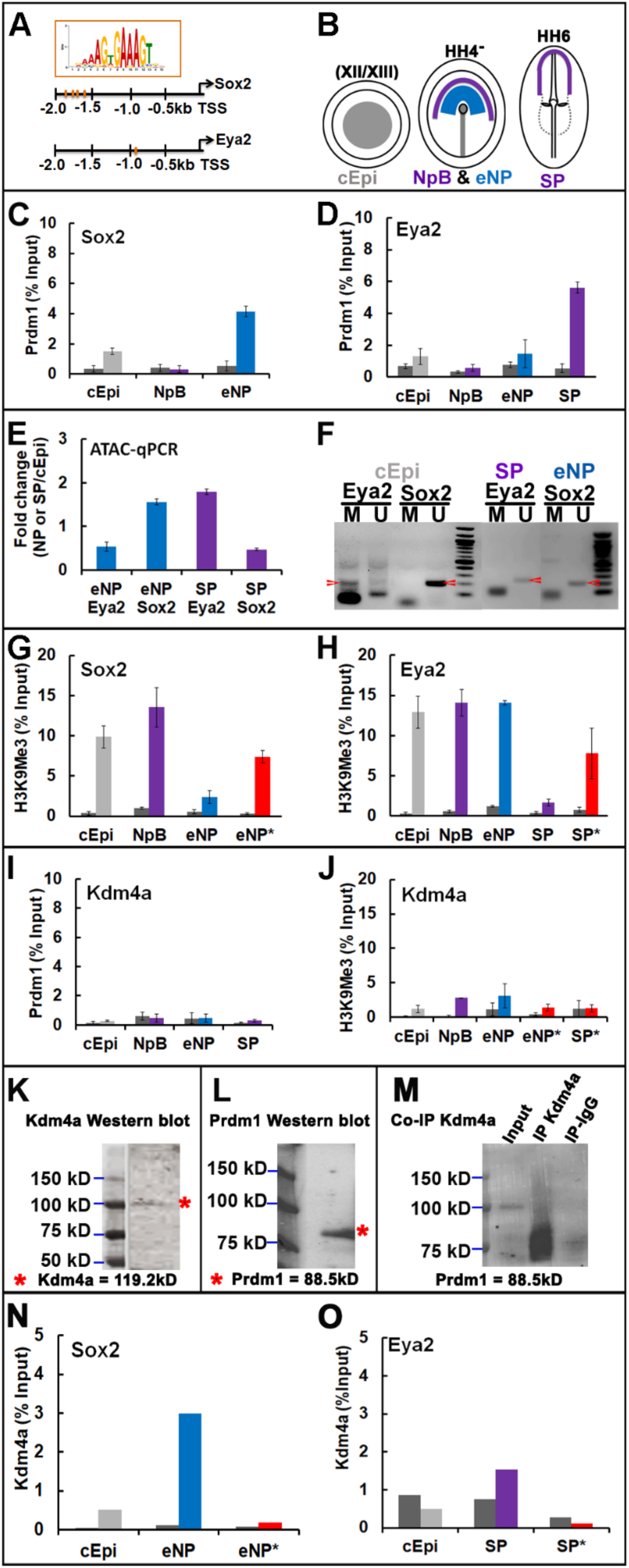
PRDM1 recruits Kdm4a to the TSS of Sox2 and Eya2 to remove repressive marks in a time-specific manner. **A.** PRDM1 binding motifs are detected upstream of the TSS of Sox2 and Eya2 (red bars). **B.** Pre-primitive streak stage epiblast (cEpi, grey), early neural plate (eNP, blue), neural plate border (NpB purple) and sensory progenitors (SP, purple) were dissected from chick embryos at different stages. **C.** Chromatin isolated from early epiblast (cEpi) neural plate border (NpB) and early neural plate (eNP) was subjected to ChIP with IgG controls (dark grey bars) and PRDM1 antibodies followed by qPCR using primers flanking the PRDM1 motifs upstream of the Sox2 TSS. PRDM1 binds to the Sox2 promoter in eNP cells. Error bars represent standard deviation from experiments carried out in triplicates on three independent occasions. **D.** Chromatin isolated from early epiblast (cEpi) neural plate border (NpB), early neural plate (eNP) and sensory progenitors (SP) was subjected to ChIP with IgG controls (dark grey bars) and PRDM1 antibodies followed by qPCR using primers flanking the PRDM1 motifs upstream of the Eya2 TSS. PRDM1 binds to the Eya2 promoter in SPs. Error bars represent standard deviation from experiments carried out in triplicates on three independent occasions. **E.** ATAC-qPCR amplifying the region containing the PRDM1 motif upstream of the Sox2 and Eya2 TSS, respectively, from eNP (blue) and SPs (purple). Bar diagrams show the fold change relative to ATAC-qPCR of the same region from pre-streak epiblast cells. The promoter region of Sox2 is accessible in neural plate, but not sensory progenitor cells, while the opposite is true for Eya2. **F.** DNA methylation upstream of the Sox2 and Eya2 TSS was assessed by bisulfide conversion of non-methylated cytosine to uracil followed by PCR with primers for specific for methylated and non-methylated DNA. At pre-streak stages (cEpi), the Eya2 TSS is largely methylated (red arrowhead), but non-methylated in SPs (red arrowhead), while the Sox2 TSS is non-methylated in epiblast and early neural plate cells (red arrowhead). **G.** ChIP-qPCR with control IgG (dark grey bars) and H3K9me3 antibodies (coloured bars) show the presence of repressive marks at the Sox2 TSS in cEpi (grey) and NpB cells (purple). H3K9me3 is reduced in the early neural plate (eNP, blue), but increases when PRDM1 is knocked down (*, red bar). **H.** ChIP-qPCR with control IgG (dark grey bars) and H3K9me3 antibodies (coloured bars) show the presence of repressive marks at the Eya2 TSS in cEpi (grey), NpB (purple) and eNP cells (blue). H3K9me3 is reduced in the sensory progenitors (SP, purple), but increases when PRDM1 is knocked down (*, red bar). **I.** PRDM1 ChIP qPCR for the region upstream of the Kdm4a TSS in different tissues shows the absence of PRDM1 binding. Dark grey bars: IgG control; coloured bars: PRDM1 antibody. J. H3K9me3 ChIP qPCR for the Kdm4a TSS in different tissues shows the absence of repressive marks and no changes over time. Dark grey bars: control IgG; coloured bars: H3K9me3 antibody. Knock-down of PRDM1 in eNP (*) and SP (*) has no effect on H3K9me3 deposition (red bars). **K.** Western blot on protein lysates from mixed eNP/SP cells with Kdm4a antibodies. **L.** Western blot on protein lysates from mixed eNP/SP cells with PRDM1 antibodies. **M.** Immunoprecipitation with Kdm4a antibodies followed by Western blot with PRDM1 antibodies. N. Kdm4a ChIP qPCR for the Sox2 TSS from epiblast (cEpi) and early neural plate (eNP) reveals binding of Kdm4a in eNP. Dark grey bars: IgG control; coloured bars: Kdm4a. Knockdown of PRDM1 in eNP (*) abolishes Kdm4a binding (red bar).

### Chromatin accessibility, DNA methylation and repressive histone mark H3K9me3 occupancy dynamically regulate *Sox2* and *Eya2* in vivo

Although *PRDM1* is expressed in all four cell populations tested, it binds close to the Sox2 and Eya2 TSS in a stage-and tissue-specific manner. We reasoned that changes in chromatin accessibility, DNA methylation and/or repressive histone marks may be important for the regulation of neural and sensory progenitor gene expression and for PRDM1 binding. To assess these features as cells become successively specified we dissected pre-streak epiblast, neural plate and sensory progenitor cells. To investigate chromatin accessibility we performed ATAC-qPCR probing the genomic regions upstream of the Sox2 and Eya2 TSS that harbour PRDM1 motifs and determined accessibility in neural plate and sensory progenitors relative to pre-streak epiblast cells. This analysis reveals that the PRDM1 site upstream of the Sox2 TSS opens in neural plate cells at HH4^−^, while that upstream of the Eya2 TSS becomes accessible only in sensory progenitors at HH5/6 (Fig. 3E). Thus, changes in chromatin accessibility coincide with the onset of *Sox2* and *Eya2* expression and with PRDM1 occupancy.

Methylation of CpG islands in proximity of the TSS is generally associated with transcriptional repression, and PRDM1 binding is known to be methylation sensitive (Doody et al., 2010). Indeed, CpG islands are predicted at position –2kb to −0.4kb from the Sox2 and –2kb to −1.7kb from the Eya2 TSS close to PRDM1 binding sites. To investigate CpG methylation we performed bisulphite assays, where cytosine is converted into uracil, while 5-methylcytosine remains intact, in pre-streak epiblast, neural plate and sensory progenitor cells. The above genomic regions were probed by PCR using primers specific for methylated and non-methylated DNA. We find that at pre-streak stages, Sox2 is not methylated, while CpG islands upstream of the Eya2 TSS are, but their methylation is lost in sensory progenitors (Fig. 3F). Thus, both genes are differentially prepared for transcription in agreement with their onset of expression at primitive streak (*Sox2*) and neural plate stages (*Eya2*).

Histone3 lysine9 trimethylation (H3K9me3) is a hallmark of transcriptional silencing and is enriched at repressed and bivalent promoter regions. To assess histone methylation we performed ChIP using H3K9me3 antibodies revealing dynamic occupancy close to the Sox2 and Eya2 TSSs. In pluripotent epiblast and HH4^−^ neural plate border cells H3K9me3 is enriched at the TSS of both genes. While H3K9me3 occupancy is reduced upstream of the Sox2 TSS in the HH4^−^ neural plate, the Eya2 TSS loses H3K9me3 later in HH5/6 sensory progenitors (Fig. 3G, H). To ensure specificity we used the ubiquitously expressed *Kdm4a* as control; we do not observe any changes in H3K9me3 (Fig 3J). Thus, histone demethylation at the TSS of Sox2 and Eya2 clearly reflects the time and tissue specific transcriptional status of both genes.

Together, these results show dynamic epigenetic changes in proximity of the TSS of the neural plate specifier Sox2 and the sensory progenitor specifier Eya2 consistent with the onset of their transcription and PRDM1 binding. In pre-streak epiblast cells, the genomic regions close to the TSS of both genes are closed, decorated by repressive H3K9me3 marks and PRDM1 does not bind. While CpG islands close to the Eya2 TSS are methylated at pre-streak stages, those proximal to Sox2 are not, foreshadowing its expression at primitive streak stages. H3K9me3 marks are lost concomitant with PRDM1 binding suggesting that PRDM1 plays an active role in preparing the onset of *Sox2* and *Eya2* transcription.

### PRDM1 recruits Kdm4a to remove repressive histone marks

PRDM1 is known to form multi-protein complexes that are generally involved in transcriptional repression. Our results suggest however that during neural and sensory progenitor specification, PRDM1 plays an activating role. To assess whether PRDM1 is required for histone demethylation upstream of the Sox2 and Eya2 TSSs, we electroporated PRDM1 aONs into neural and sensory progenitors at HH3 and collected early neural plate and sensory progenitors after 16-24 hrs. ChIP-qPCR reveals that H3K9 trimethylation increases in the absence of PRDM1 when compared to controls (Fig. 3G, H). In contrast, there is no change in H3K9me3 at the Kdm4a TSS (Fig. 3J). These observations lead us to propose that PRDM1 mediates activation of *Sox2* and *Eya2* transcription by promoting the removal of repressive histone marks.

PRDM1 itself does not have any enzymatic activity but is known to interact with different chromatin modifiers (Ancelin et al., 2006; Gyory et al., 2004; Magnusdottir et al., 2013; for review: Bikoff et al., 2009; Mzoughi et al., 2016). The histone demethylase Kdm4a specifically removes trimethylation from H3K9 and is broadly expressed in the chick epiblast (Strobl-Mazzulla et al., 2010). We therefore first assessed whether PRDM1 and Kdm4a interact. Western blot analysis from mixed neural plate border and sensory progenitors confirms the expression of both proteins (Fig. 3K, L), while co-immunoprecipitation reveals that PRDM1 and Kdm4a bind to each other (Fig. 3M). We next examined whether Kdm4a is located close to the Sox2 and Eya2 TSS using ChIP qPCR. In pre-streak epiblast, Kdm4a does not occupy either region, but binds close to the Sox2 TSS in early neural plate cells and close to the Eya2 TSS in sensory progenitor cells (Fig. 3N, O). Thus, both PRDM1 and Kdm4a are found close to the TSS of Sox2 and Eya2 when their transcription becomes activated.

Is PRDM1 required for Kdm4a binding? To test this, we knocked down PRDM1 in neural and sensory progenitors by electroporation of aONs at HH3 and collected both tissues at HH4^−^ and HH6, respectively. Kdm4a ChIP-qPRC reveals that in the absence of PRDM1, Kdm4a binding to both genomic regions is lost (Fig. 3N, O). These results show that PRDM1 is required to recruit Kdm4a to the TSS of Sox2 and Eya2.

Together our results provide a model for sequential specification of neural and sensory progenitor fates in the embryonic ectoderm, with PRDM1 as a central player (Fig. 4). Prior to activation of cell type specific transcripts their TSSs are inaccessible and decorated with repressive marks like H3K9me3. As they become accessible, PRDM1 binds upstream of the TSS of key neural and sensory progenitor genes, recruits the histone demethylase Kdm4a, which in turn demethylates H3K9me3 to facilitate transcriptional activation.

**Figure 4.**
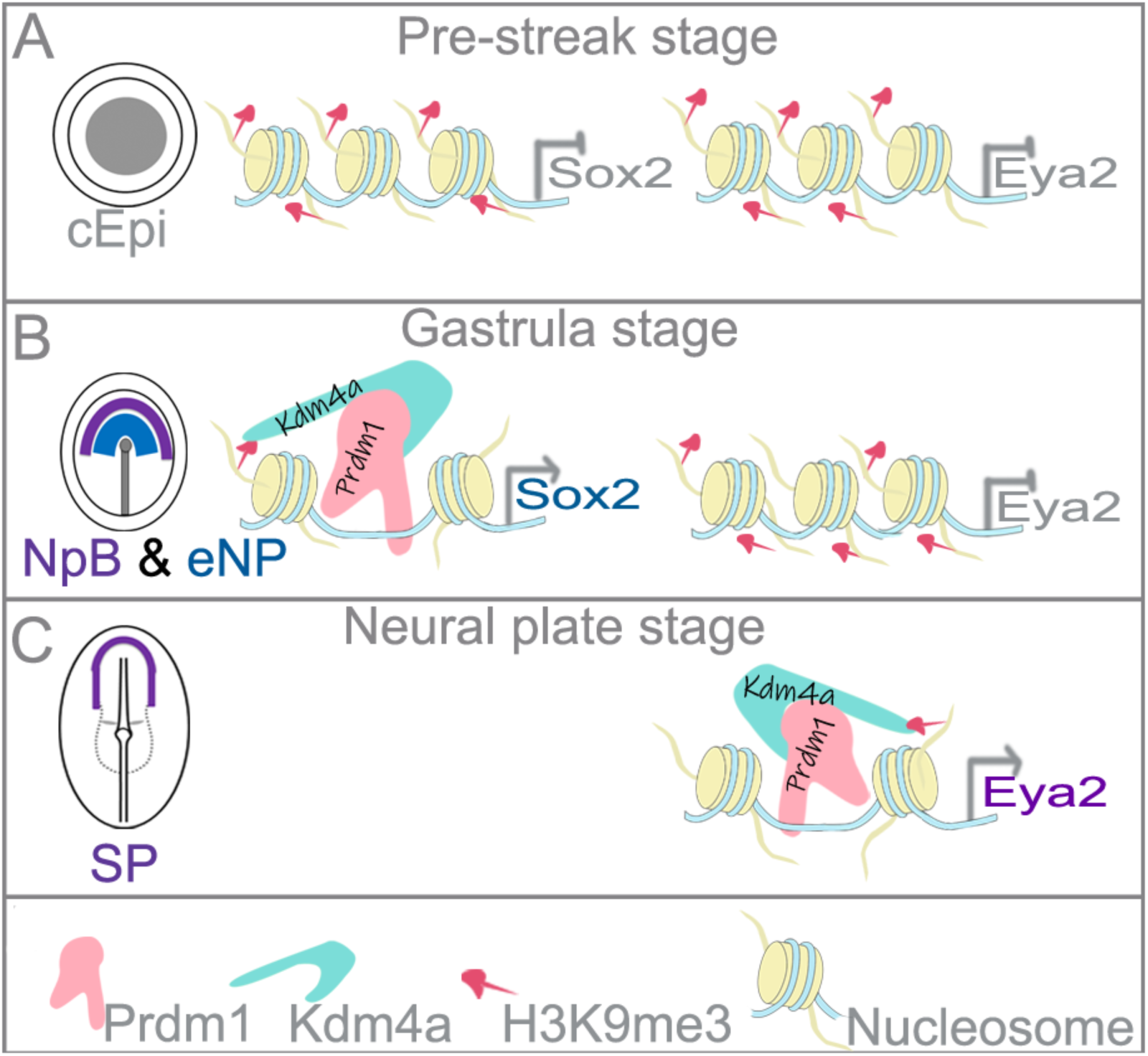
Model for PRDM1 function regulating *Sox2* and *Eya2*. **A.** At pre-primitive streak stages, the promoter region of Sox2 and Eya2 is closed and occupied by repressive H3K9me3 marks. **B.** At early gastrula stages, *PRDM1* is expressed broadly in the ectoderm. The Sox2 promoter becomes accessible and PRDM1 binds upstream of the TSS recruiting the demethylase Kdm4a, which removes repressive H3K9me3 to allow *Sox2* transcription. **C.** At neural plate stages, the Eya2 promoter region opens in sensory progenitors, allowing PRDM1 binding and recruitment of Kdm4a. *Eya2* begins to be transcribed.

## DISCUSSION

During embryo development, exit from pluripotency and sequential activation of distinct differentiation programmes must tightly controlled to coordinate cell fate decisions with morphogenetic processes. Our findings place the transcription factor PRDM1 into the centre of the network regulating these processes in the embryonic ectoderm. Dissection of PRDM1 function in time and space allows us to distinguish different PRDM1 activities. Initially, PRDM1 is required for cells to lose pluripotency markers, and at the same time for activating the neural, neural crest and sensory progenitor programme. Once cells are specified PRDM1 must be downregulated rapidly to maintain these fates: prolonged expression prevents the differentiation of neural lineages. Thus, PRDM1 function changes rapidly presumably due to interaction with different co-factors.

### PRDM1 balances loss of pluripotency markers and the activation of neural programmes

In amniote embryonic stem cells, different epigenetic mechanisms including transcriptional repressors, histone and DNA methylation maintain pluripotency, while simultaneously preventing premature expression of differentiation markers (Andrey and Mundlos, 2017; Habibi and Stunnenberg, 2017; Kalkan and Smith, 2014; Kim et al., 2008; Li and Izpisua Belmonte, 2018; Schlesinger and Meshorer, 2019; Surani et al., 2007; Theunissen and Jaenisch, 2017). In the embryo, the loss of pluripotency is tightly coordinated with morphogenetic events. As the three germ layers form, cells in the ectoderm are rapidly specified as central and peripheral nervous system progenitors while pluripotency gene expression decreases. In chick, the pluripotency associated factors *PouV/Oct4, Nanog, Sox3* and *ERNI/Ens1* are expressed in the blastoderm prior to primitive streak formation (Jean et al., 2015; Lavial et al., 2007; Streit et al., 2000), as is *PRDM1* (this study). Like *ERNI, PRDM1* expression is lost in epiblast cells as differentiation programmes are activated. Our results suggest that PRDM1 plays a dual role: while *PRDM1* knock-down leads to upregulation of pluripotency markers, neural, neural crest and sensory progenitor specifiers fail to be expressed. Interestingly, a similar scenario has been observed in primordial germ cells and during their conversion into pluripotent cells (Nagamatsu et al., 2015; Surani et al., 2007). PRDM1 deletion in primordial germ cells enhances their dedifferentiation into pluripotent embryonic germ cells, paralleling our findings for neural fates in the ectoderm. Conversely, its overexpression in embryonic stem cells suppresses parts of the pluripotency network and prevents the conversion of *in-vitro* induced primordial germ cells into pluripotent embryonic germ cells. Together, these findings highlight PRDM1 as an important node in the network that controls the balance between pluripotency and differentiation for several different lineages.

### Molecular events controlling neural, neural crest and sensory progenitor fates

Around the time of gastrulation ectodermal cells begin to activate neural, neural crest and sensory progenitor genes in a temporal sequence. Here we demonstrate that prior to this, the TSSs of the neural marker *Sox2* and the sensory progenitor marker *Eya2* are not accessible, are decorated with the repressive histone marks H3K9me3 and not bound by PRDM1. H3K9me3 is linked to gene silencing and known to bind the transcriptional repressors HP1 (Bannister et al., 2001; Lachner et al., 2001; Nielsen et al., 2001), and this is likely to prevent inappropriate transcription of both genes. As development proceeds, both promoter regions open in a time-and tissue-specific manner allowing PRDM1 to bind and recruit the demethylase Kdm4a leading to removal of repressive H3K9me3. Previous studies have shown that at pre-streak stages, the N2 enhancer of Sox2 is occupied by a complex including Geminin and ERNI, which in turn recruits HP1γ to repress *Sox2* transcription (Papanayotou et al., 2008). During gastrulation, the coiled-coil protein *BERT* displaces ERNI from the complex together with HP1γ. Here we show that at the same time PRDM1 enables the removal of repressive marks at the promoter region suggesting that both mechanisms work in concert to activate *Sox2* transcription in early neural plate cells. It is tempting to speculate that a similar mechanism acts to promote the transcription of neural plate border, neural crest and sensory progenitor genes. Our analysis shows that the neural plate border genes Dlx5/6, Gata3, TFAPα/ε and Msx2 harbour PRDM1 motifs close to their TSS, as do Six1 and Eya2, and the neural crest factor Foxd3, suggesting that PRDM1 may indeed control the onset of their expression directly recruiting Kdm4a to remove repressive histone marks. Whether Geminin, ERNI and BERT at enhancer regions cooperate with PRDM1 as described for Sox2 remains to be elucidated. Together, these observations place PRDM1 into the centre of the transcriptional network controlling the onset of neural, neural crest and sensory progenitor specification.

### A functional switch of PRDM1: from activator to repressor over time

PRDM1 is generally considered to act as a transcriptional repressor. Here, we provide evidence that in early epiblast cells it functions as a transcriptional activator and elucidate the underlying mechanism. PRDM1 contains a proline-serine rich domain and five C2H2 zing fingers; the latter is responsible for DNA binding, while both are involved in the recruitment of additional cofactors (Ancelin et al., 2006; Gyory et al., 2004; Kurimoto et al., 2015; Ren et al., 1999; for review: Bikoff et al., 2009; Mzoughi et al., 2016). In primordial germ cells, PRDM1 is required for the repression of somatic genes and forms a complex with the arginine-specific histone methyltransferase Prmt5, which in turn mediates methylation of histone H2A and H4 tails (Ancelin et al., 2006). In B cells, it represses genes associated with cell cycle progression and B cell maturation like *c-Myc, CIITA* and *Pax5* by recruiting histone deacetylases to their promoters (Bikoff et al., 2009; Gyory et al., 2004; Lin et al., 1997; Yu et al., 2000). In addition, interacting with Groucho proteins or the G9a methyltransferase it represses the expression of interferon-β (Gyory et al., 2004; Ren et al., 1999).

In contrast, our results reveal that PRDM1 acts as an activator in neural, neural crest and sensory progenitor cells. As in chick, zebrafish PRDM1 is required for neural crest cell formation by interacting with the enhancers of the neural crest factors *Foxd3* and *Tfap2a* (Powell et al., 2013). Our results show that PRDM1 physically interacts with the histone demethylase Kdm4a and recruits near the TSS of the neural specifier Sox2 and the sensory progenitor specifier Eya2, which in turn leads to reduced H3K9me3 occupancy and gene activation. Both Kdm4a binding and loss of the repressive mark H3K9me3 are PRDM1 dependent, explaining why PRDM1 is necessary for the onset of expression of both genes. While we do not provide evidence for direct Foxd3 activation by PRDM1, it is likely that a similar mechanism operates in neural crest cells. In *Xenopus*, Kdm4a overexpression leads to upregulation of *Foxd3* and the neural crest gene *Slug1* accompanied by a loss of H3K9me3 at the Foxd3 promoter (Powell et al., 2013). In chick, Kdm4a is required for neural crest cell development and mediates H3K9me3 demethylation close to the TSS of the neural crest specifiers Sox10 and Snail2 (Strobl-Mazzulla et al., 2010). We suggest that in this scenario PRDM1 recruits Kdm4a to the promoters of Foxd3, Sox10 and Snail2 and is therefore crucial for their activation (Matsukawa et al., 2015). Interestingly, in sensory progenitor cells the promoters of neural crest cell specifiers are occupied by PRDM12, which represses their expression by promoting H3K9me3 deposition and thus prevents inappropriate expression of neural crest genes.

While promoting transcriptional activation of neural determinants in early epiblast cells, *PRDM1* switches its function to repress neural lineages shortly thereafter. During normal development *PRDM1* is rapidly downregulated as lineage-specific genes become expressed, and we show that the fate specifiers themselves play a crucial role: misexpression of Sox2, Foxd3 or Six1 leads to loss of *PRDM1*. Likewise, in zebrafish Foxd3 represses *PRDM1* (Powell et al., 2013). In contrast, prolonging *PRDM1* expression inhibits neural, neural crest and sensory progenitor fate. It is likely that in this context PRDM1 recruits transcriptional repressors like histone deacetylases or Groucho family members. Thus, at early neural plate stages PRDM1 and Sox2, Foxd3 and Six1 mutually repress each other and the loss of PRDM1 after cell fate specification allows neural, neural crest and sensory progenitor cells to maintain their identity. A tight regulation of Prdm family members and their interacting partners is therefore required for fine tuning gene expression at the neural plate border and for mediating cell fate choices mediating.

## CONCLUSION

During embryo development, exit from pluripotency and sequential activation of distinct differentiation programmes must tightly controlled in time and space to coordinate cell fate decisions with morphogenetic processes. PRDM1 emerges as a key node in the network regulating these processes in the embryonic ectoderm (Fig. 4). PRDM1 is required for cells to lose pluripotency markers, while at the same time activating the neural, neural crest and sensory progenitor programme. Once initiated PRDM1 must be downregulated rapidly to allow central and peripheral nervous system precursors to maintain their identity. Thus, PRDM1 is a versatile factor which works both as transcriptional activator or repressor in a time dependent manner to control fate decisions. In embryonic stem cells and primordial germ cells, PRDM1 downstream targets have been extensively characterised and are partially overlapping. It will be interesting to evaluate how these networks diverge in the neural lineage.

## MATERIALS AND METHODS

### Embryo collection and whole mount *in situ* hybridization

Fertile hens’ eggs were obtained from Henry Stewart farms and incubated at 38°C until they reached the stage required (Hamburger and Hamilton, 1951). Embryos were collected in nuclease-free phosphate buffered saline (PSB) and fixed in 4% paraformaldehyde at room temperature for 4-5 hours. Whole-mount *in situ* hybridization was carried out as previously described (Streit and Stern, 2001). To generate antisense Digoxigenin labelled probes the following plasmids were used: PRDM1 (Chen et al., 2017), Six1 (Sato et al., 2010), Eya2 (Mishima and Tomarev, 1998), Dlx5 (McLarren et al., 2003), Sox2 (Rex et al., 1997), Foxd3 (Kos et al., 2001), ERNI (Streit et al., 2000) and Gata3 (Sheng and Stern, 1999).

### Electroporation of antisense oligonucleotides and expression vectors

Primitive streak stage embryos were electroporated in Tyrode’s saline using five pulses of 5-7mV for 50ms with an interval of 750ms and cultured in modified New culture (Stern and Ireland, 1981) until the 1-5 somite stages. Two splicing-blocking antisense oligonucleotides (aONs) were designed to knock down *PRDM1* by pre-mRNA mis-splicing: aON1 (5’-ACTGTAATGCACTTACTGAGGTTC-3’) targets the exon6-intron6 and aON2 (5’-TCTTAGTCTCCACCACCTAC-CTTCA-3’) targets exon7-intron7 boundary. Control ONs were 5’-CCTCTTACCTCAGTTACAATTTATA-3’ (GeneTools). For electroporation, each ON was used at a final concentration of 1mM in distilled water containing 6% Sucrose, 0.04% Fast Green and 0.5mg/ml carrier plasmid (puc19). All ONs were labelled with fluorescein; to visualise targeted cells we performed immunocytochemistry using anti-fluorescein antibodies (Roche 426346910).

For misexpression, the coding regions of PRDM1, Six1, Foxd3 and Sox2 (gift from C. D. Stern) were cloned into pCAB-IRES-eGFP vectors, which drives ubiquitous expression of the gene of interest and eGFP. For electroporation plasmids were used at a concentration of 2mg/ml in distilled water containing 6% Sucrose and 0.04% Fast Green. Anti-GFP antibodies (Life Technology, a11122) and Alexa 488 coupled secondary antibodies were used to visualize targeted cells.

### NanoString nCounter

HH3^+^/4^−^ embryos were electroporated with aONs targeting PRDM1 or control ONs, allowed to grow until HH6 and targeted sensory progenitor cells were dissected using a fluorescence microscope. Each sample contained 5-10 tissue pieces (5000-7000 cells), which were immediately lysed in lysis buffer and processed for NanoString nCounter as previously described (Hintze et al., 2017). Each experiment was repeated on three independent occasions. Counts were normalized to the positive controls contained within the hybridization mix and negative control probe values were used to determine the background threshold level. Transcripts with expression values below the threshold were removed from further analysis. Counts were then normalized to the total amount of mRNA in each sample and differential expression between control and experimental conditions was determined using an unpaired two-tailed Student’s *t*-test comparing the average of three biological replicates (p<0.05, > 1.2-fold change).

### Chromatin immunoprecipitation

For chromatin immunoprecipitation (ChIP) 15-20 explants of pre-streak epiblast, HH4^−^ neural plate border and early neural plate, and HH6 sensory progenitors were dissected in Tyrode’s saline. Tissues were homogenized in nuclear extraction buffer (NEB: 0.5% NP-40, 0.25% TritonX-100, 10mM Tris-HCl pH7.5, 3mM CaCl2, 0.25M Sucrose, 1mM DDT, 0.2mM PMSF, 1× Protease Inhibitor (PI)) using a Dounce homogenizer and fixed with 0.9% formaldehyde for 10 min at room temperature. The fixing reaction was quenched with 125mM glycine, the tissues were washed three times in PSB-PI (1mM DDT, 0.2mM PMSF, 1× PIS). Cells were re-suspended in NEB and nuclei were released by Dounce homogenizing using a tight pestle. Nuclei were washed with PSB-PI and lysed in SDS lysis buffer (1% SDS, 10mM EDTA in 50mM Tris-HCl pH8) for 1 hour on ice before being diluted to 0.9ml with ChIP Dilution Buffer (CDB: 0.01%SDS, 1.2mM EDTA, 167mM NaCl, 1mM DDT, 0.2mM PMSF and 1× PI in 16.7mM Tris-HCl, pH8) and sonicated to obtain 500-1000 bp chromatin fragments. TritonX-100 was added to a final concentration of 1% before the chromatin was used for ChIP. Protein-A magnetic beads (100ul) were blocked with 0.5% BSA and coated with the 5mg antibody (Abcam; PRDM1 ab13700; H3K9me3 ab8898; control IgG 5mg) before being added to the chromatin and allowed to bind overnight at 4°C. Beads were washed six times with RIPA buffer (500mM LiCl, 1mM EDTA, 1%NP-40, 0.7% Na-deoxycholate, 1× PI in 50mM HEPES-KOH, pH8), followed by three washes in 10mM Tris-HCl, pH8 containing 1mM EDTA and 50mM NaCl. Chromatin was released from the beads in elution buffer (10mM EDTA, 1%SDS in 50mM Tris-HCl, pH8) at 65°C for 30 minutes. The eluted chromatin was reverse cross-linked by incubating 65°C for overnight before being incubated with RNaseA (0.2mg/ml) and Proteinase K(0.2mg/ml). DNA was purified using phenol-chloroform and assayed with qPCR using primers for different genomic regions flanking PRDM1 morida (see Supplemental Table 3). The genomic region −2kb from the TSS of each gene was extracted from GalGal6 and screened for PRDM1 motifs using RSAT (rsat.sb-roscoff.fr). The PRDM1 matrix obtained from JASPER (jaspar.genereg.net).

### Western blot and co-immunoprecipitation

For western blot, 50 HH4-6 neural plate border / sensory progenitor tissues we lysed in SDS-PAGE loading buffer by heating to 100°C for 10min. The lysate was separated using 10% SDS-PAGE and proteins were transferred to immuno-blot PVDF membrane. Blots were blocked with 5% milk powder in PBS for 1 hour at room temperature, followed by incubation with primary antibodies to PRDM1 (Abcam: ab13700, 1:200) and Kdm4a (Abcam: ab24545, 1:400) at 4°C overnight. After washing in PBS, 0.2% triton-100, blots were incubated with HRP-conjugated secondary antibodies (Abcam: Donkey ant-goat-HRP, ab6885; Jackson Labs Technology: Goat ant-rabbit-HRP, 111-035-003, 1:1000) for 1h at room temperature, washed again, developed using clarity western ECL (Biorad: 170-5-60) and imaged with Biorad ChemiDoc touch imaging system.

For co-immunoprecipitation (CoIP) tissues were lysed in the CoIP buffer (100mM NaCl, 0.2% triton-100, 0.5% NP-40, 2mM β-mercaptoethanol, 1mM DTT, protease inhibitor in 20mM Tris pH7.5), incubated with Kdm4a antibody (5mg) overnight at 4°C. Kdm4a antibody bound proteins were precipitated using Protein-A coated Dynabeads(100μl). After three washes with CoIP-Buffer bead were suspended in SDS-PAGE loading buffer and proteins were process for Western blot.

### Assay for Transposase-Accessible Chromatin qPCR (ATAC pPCR)

To assess the accessibility of the chromatin upstream of the Sox2 and Eya2 TSSs we performed ATAC qPCR from sensory progenitors, neural plate and pre-streak epiblast. 15 sensory progenitor and neural plate explants (HH6) and two central pre-streak epiblast pieces (about 10,000 cells each) were used for each experiment. Tissues were dissociated and nuclei were isolated in cold lysis buffer (10mM NaCl, 3mM Mgcl2, 0.1% IPGEPAL CA-630 in 10mM Tris-HCL, pH7.4). Nuclei were washed with lysis buffer, recovered by centrifugation at 500xg and treated with transposase (Tn5 Transposase, Illumina, FC-121-1030) for 10min as described by Buenrostro and colleagues (Buenrostro et al., 2013). DNA was purified using a mini-elute PCR purification kit (QIAGEN) and qPCR was performed using primers for the region upstream of the Sox2 and Eya2 TSS.

### DNA methylation assay

To examine the methylation status CpG islands were predicted within 2kb upstream of upstream of the Sox2 and Eya2 TSS using MethPrimer (Li and Dahiya, 2002). Central pre-streak epiblast, early neural plate (HH4^−^) and sensory progenitors (HH6) were dissected and genomic DNA was prepared using the DNeasy blood & tissue kit from Qiagen (69504). Genomic DNA was treated with bisulfide to convert unmethylated cytosine to uracil using Thermo Scientific EpiJET Bisulfide Conversion Kit (Fisher Scientific, K1461) and used for PCR using two pairs of primers for the region upstream the TSS of Sox2 and Eya2 that that flank the PRDM1 motif and amplify either methylated or unmethylated DNA. PCR was carried out using Phusion U Hot Start DNA Polymerase (Fisher Scientific F-555S/L) and analysed by gel electrophoresis.

## ACKNOWLEDGEMENTS

The authors are grateful to Ewa Kolano for excellent technical assistance, to Claudio Stern and Karen Liu for critical reading of the manuscript, and the Streit group and Jeremy Green for many discussions. This work was funded by project grants to A.S. from the National Institutes of Health (DE022065; DC011577) and the Biotechnology and Biological Sciences Research Council (BB/I021647/1).

## Author contributions

A.S. together with R.S.P. designed the project. R.S.P. and M.H. conducted the embryo experiments; R.S.P. carried out the molecular experiments and prepared the figures. A.S. supervised and administered the project, obtained the funding and wrote the manuscript.

## REFERENCES

Ancelin, K., Lange, U.C., Hajkova, P., Schneider, R., Bannister, A.J., Kouzarides, T., Surani, M.A., 2006. Blimp1 associates with Prmt5 and directs histone arginine methylation in mouse germ cells. Nat Cell Biol 8, 623–630.

Andrey, G., Mundlos, S., 2017. The three-dimensional genome: regulating gene expression during pluripotency and development. Development 144, 3646–3658.

Bannister, A.J., Zegerman, P., Partridge, J.F., Miska, E.A., Thomas, J.O., Allshire, R.C., Kouzarides, T., 2001. Selective recognition of methylated lysine 9 on histone H3 by the HP1 chromo domain. Nature 410, 120–124.

Basch, M.L., Bronner-Fraser, M., Garcia-Castro, M.I., 2006. Specification of the neural crest occurs during gastrulation and requires Pax7. Nature 441, 218–222.

Bikoff, E.K., Morgan, M.A., Robertson, E.J., 2009. An expanding job description for Blimp-1/PRDM1. Curr Opin Genet Dev 19, 379–385.

Buenrostro, J.D., Giresi, P.G., Zaba, L.C., Chang, H.Y., Greenleaf, W.J., 2013. Transposition of native chromatin for fast and sensitive epigenomic profiling of open chromatin, DNA-binding proteins and nucleosome position. Nat Methods 10, 1213–1218.

Buitrago-Delgado, E., Nordin, K., Rao, A., Geary, L., LaBonne, C., 2015. Shared regulatory programs suggest retention of blastula-stage potential in neural crest cells. Science 348, 1332–1335.

Buitrago-Delgado, E., Schock, E.N., Nordin, K., LaBonne, C., 2018. A transition from SoxB1 to SoxE transcription factors is essential for progression from pluripotent blastula cells to neural crest cells. Dev Biol 444, 50–61.

Chen, J., Tambalo, M., Barembaum, M., Ranganathan, R., Simoes-Costa, M., Bronner, M.E., Streit, A., 2017. A systems-level approach reveals new gene regulatory modules in the developing ear. Development 144, 1531–1543.

Doody, G.M., Care, M.A., Burgoyne, N.J., Bradford, J.R., Bota, M., Bonifer, C., Westhead, D.R., Tooze, R.M., 2010. An extended set of PRDM1/BLIMP1 target genes links binding motif type to dynamic repression. Nucleic Acids Res 38, 5336–5350.

Dunn, S.J., Martello, G., Yordanov, B., Emmott, S., Smith, A.G., 2014. Defining an essential transcription factor program for naive pluripotency. Science 344, 1156–1160.

Ezin, A.M., Fraser, S.E., Bronner-Fraser, M., 2009. Fate map and morphogenesis of presumptive neural crest and dorsal neural tube. Dev Biol 330, 221–236.

Fernandez-Garre, P., Rodriguez-Gallardo, L., Gallego-Diaz, V., Alvarez, I.S., Puelles, L., 2002. Fate map of the chicken neural plate at stage 4. Development 129, 2807–2822.

Fernandez-Tresguerres, B., Canon, S., Rayon, T., Pernaute, B., Crespo, M., Torroja, C., Manzanares, M., 2010. Evolution of the mammalian embryonic pluripotency gene regulatory network. Proc Natl Acad Sci U S A 107, 19955–19960.

Fujita, K., Ogawa, R., Ito, K., 2016. CHD7, Oct3/4, Sox2, and Nanog control FoxD3 expression during mouse neural crest-derived stem cell formation. FEBS J 283, 3791–3806.

Gyory, I., Wu, J., Fejer, G., Seto, E., Wright, K.L., 2004. PRDI-BF1 recruits the histone H3 methyltransferase G9a in transcriptional silencing. Nat Immunol 5, 299–308.

Habibi, E., Stunnenberg, H.G., 2017. Transcriptional and epigenetic control in mouse pluripotency: lessons from in vivo and in vitro studies. Curr Opin Genet Dev 46, 114–122.

Hamburger, V., Hamilton, H.L., 1951. A series of normal stages in the development of the chick embryo. J Morph 88, 49–92.

Hernandez-Lagunas, L., Choi, I.F., Kaji, T., Simpson, P., Hershey, C., Zhou, Y., Zon, L., Mercola, M., Artinger, K.B., 2005. Zebrafish narrowminded disrupts the transcription factor prdm1 and is required for neural crest and sensory neuron specification. Dev Biol 278, 347–357.

Hintze, M., Prajapati, R.S., Tambalo, M., Christophorou, N.A.D., Anwar, M., Grocott, T., Streit, A., 2017. Cell interactions, signals and transcriptional hierarchy governing placode progenitor induction. Development 144, 2810–2823.

Jean, C., Oliveira, N.M., Intarapat, S., Fuet, A., Mazoyer, C., De Almeida, I., Trevers, K., Boast, S., Aubel, P., Bertocchini, F., Stern, C.D., Pain, B., 2015. Transcriptome analysis of chicken ES, blastodermal and germ cells reveals that chick ES cells are equivalent to mouse ES cells rather than EpiSC. Stem cell research 14, 54–67.

Kalkan, T., Smith, A., 2014. Mapping the route from naive pluripotency to lineage specification. Philosophical Transactions of the Royal Society B: Biological Sciences 369, 20130540.

Kallies, A., Nutt, S.L., 2007. Terminal differentiation of lymphocytes depends on Blimp-1. Curr Opin Immunol 19, 156–162.

Khudyakov, J., Bronner-Fraser, M., 2009. Comprehensive spatiotemporal analysis of early chick neural crest network genes. Dev Dyn 238, 716–723.

Kim, J., Chu, J., Shen, X., Wang, J., Orkin, S.H., 2008. An extended transcriptional network for pluripotency of embryonic stem cells. Cell 132, 1049–1061.

Kos, R., Reedy, M.V., Johnson, R.L., Erickson, C.A., 2001. The winged-helix transcription factor FoxD3 is important for establishing the neural crest lineage and repressing melanogenesis in avian embryos. Development 128, 1467–1479.

Kurimoto, K., Yabuta, Y., Hayashi, K., Ohta, H., Kiyonari, H., Mitani, T., Moritoki, Y., Kohri, K., Kimura, H., Yamamoto, T., Katou, Y., Shirahige, K., Saitou, M., 2015. Quantitative Dynamics of Chromatin Remodeling during Germ Cell Specification from Mouse Embryonic Stem Cells. Cell Stem Cell 16, 517– 532.

Kwon, H.J., Bhat, N., Sweet, E.M., Cornell, R.A., Riley, B.B., 2010. Identification of early requirements for preplacodal ectoderm and sensory organ development. PLoS Genet 6.

Lachner, M., O’Carroll, D., Rea, S., Mechtler, K., Jenuwein, T., 2001. Methylation of histone H3 lysine 9 creates a binding site for HP1 proteins. Nature 410, 116–120.

Lavial, F., Acloque, H., Bertocchini, F., Macleod, D.J., Boast, S., Bachelard, E., Montillet, G., Thenot, S., Sang, H.M., Stern, C.D., Samarut, J., Pain, B., 2007. The Oct4 homologue PouV and Nanog regulate pluripotency in chicken embryonic stem cells. Development 134, 3549–3563.

Li, L.C., Dahiya, R., 2002. MethPrimer: designing primers for methylation PCRs. Bioinformatics 18, 1427–1431.

Li, M., Izpisua Belmonte, J.C., 2018. Deconstructing the pluripotency gene regulatory network. Nature Cell Biology 20, 382–392.

Lin, Y., Wong, K., Calame, K., 1997. Repression of c-myc transcription by Blimp-1, an inducer of terminal B cell differentiation. Science 276, 596–599.

Litsiou, A., Hanson, S., Streit, A., 2005. A balance of FGF, Wnt and BMP signalling positions the future placode territory in the head. Development 132, 4051–4062.

Lukoseviciute, M., Gavriouchkina, D., Williams, R.M., Hochgreb-Hagele, T., Senanayake, U., Chong-Morrison, V., Thongjuea, S., Repapi, E., Mead, A., Sauka-Spengler, T., 2018. From Pioneer to Repressor: Bimodal foxd3 Activity Dynamically Remodels Neural Crest Regulatory Landscape In Vivo. Dev Cell 47, 608–628 e606.

Magnusdottir, E., Dietmann, S., Murakami, K., Gunesdogan, U., Tang, F., Bao, S., Diamanti, E., Lao, K., Gottgens, B., Azim Surani, M., 2013. A tripartite transcription factor network regulates primordial germ cell specification in mice. Nat Cell Biol 15, 905–915.

Matsukawa, S., Miwata, K., Asashima, M., Michiue, T., 2015. The requirement of histone modification by PRDM12 and Kdm4a for the development of pre-placodal ectoderm and neural crest in Xenopus. Dev Biol 399, 164–176.

McLarren, K.W., Litsiou, A., Streit, A., 2003. DLX5 positions the neural crest and preplacode region at the border of the neural plate. Dev Biol 259, 34–47.

Mishima, N., Tomarev, S., 1998. Chicken Eyes absent 2 gene: isolation and expression pattern during development. Int J Dev Biol 42, 1109–1115.

Mundell, N.A., Labosky, P.A., 2011. Neural crest stem cell multipotency requires Foxd3 to maintain neural potential and repress mesenchymal fates. Development 138, 641–652.

Mzoughi, S., Tan, Y.X., Low, D., Guccione, E., 2016. The role of PRDMs in cancer: one family, two sides. Curr Opin Genet Dev 36, 83–91.

Nagamatsu, G., Saito, S., Takubo, K., Suda, T., 2015. Integrative Analysis of the Acquisition of Pluripotency in PGCs Reveals the Mutually Exclusive Roles of Blimp-1 and AKT Signaling. Stem Cell Reports 5, 111–124.

Nielsen, S.J., Schneider, R., Bauer, U.M., Bannister, A.J., Morrison, A., O’Carroll, D., Firestein, R., Cleary, M., Jenuwein, T., Herrera, R.E., Kouzarides, T., 2001. Rb targets histone H3 methylation and HP1 to promoters. Nature 412, 561–565.

Nutt, S.L., Fairfax, K.A., Kallies, A., 2007. BLIMP1 guides the fate of effector B and T cells. Nat Rev Immunol 7, 923–927.

Ohinata, Y., Payer, B., O’Carroll, D., Ancelin, K., Ono, Y., Sano, M., Barton, S.C., Obukhanych, T., Nussenzweig, M., Tarakhovsky, A., Saitou, M., Surani, M.A., 2005. Blimp1 is a critical determinant of the germ cell lineage in mice. Nature 436, 207–213.

Olesnicky, E., Hernandez-Lagunas, L., Artinger, K.B., 2010. prdm1a Regulates sox10 and islet1 in the development of neural crest and Rohon-Beard sensory neurons. Genesis 48, 656–666.

Papanayotou, C., Mey, A., Birot, A.M., Saka, Y., Boast, S., Smith, J.C., Samarut, J., Stern, C.D., 2008. A mechanism regulating the onset of Sox2 expression in the embryonic neural plate. PLoS Biol 6, e2.

Pieper, M., Ahrens, K., Rink, E., Peter, A., Schlosser, G., 2012. Differential distribution of competence for panplacodal and neural crest induction to non-neural and neural ectoderm. Development 139, 1175–1187.

Pla, P., Monsoro-Burq, A.H., 2018. The neural border: Induction, specification and maturation of the territory that generates neural crest cells. Dev Biol 444 Suppl 1, S36–S46.

Posfai, E., Tam, O.H., Rossant, J., 2014. Mechanisms of pluripotency in vivo and in vitro. Curr Top Dev Biol 107, 1–37.

Powell, D.R., Hernandez-Lagunas, L., LaMonica, K., Artinger, K.B., 2013. Prdm1a directly activates foxd3 and tfap2a during zebrafish neural crest specification. Development 140, 3445–3455.

Puelles, L., Fernandez-Garre, P., Sanchez-Arrones, L., Garcia-Calero, E., Rodriguez-Gallardo, L., 2005. Correlation of a chicken stage 4 neural plate fate map with early gene expression patterns. Brain Res Brain Res Rev 49, 167–178.

Ren, B., Chee, K.J., Kim, T.H., Maniatis, T., 1999. PRDI-BF1/Blimp-1 repression is mediated by corepressors of the Groucho family of proteins. Genes Dev 13, 125–137.

Rex, M., Orme, A., Uwanogho, D., Tointon, K., Wigmore, P.M., Sharpe, P.T., Scotting, P.J., 1997. Dynamic expression of chicken Sox2 and Sox3 genes in ectoderm induced to form neural tissue. Dev Dyn 209, 323–332.

Robertson, E.J., Charatsi, I., Joyner, C.J., Koonce, C.H., Morgan, M., Islam, A., Paterson, C., Lejsek, E., Arnold, S.J., Kallies, A., Nutt, S.L., Bikoff, E.K., 2007. Blimp1 regulates development of the posterior forelimb, caudal pharyngeal arches, heart and sensory vibrissae in mice. Development 134, 4335– 4345.

Rossant, J., Tam, P.P.L., 2017. New Insights into Early Human Development: Lessons for Stem Cell Derivation and Differentiation. Cell Stem Cell 20, 18–28.

Saitou, M., Payer, B., O’Carroll, D., Ohinata, Y., Surani, M.A., 2005. Blimp1 and the emergence of the germ line during development in the mouse. Cell Cycle 4, 1736–1740.

Sanchez-Arrones, L., Stern, C.D., Bovolenta, P., Puelles, L., 2012. Sharpening of the anterior neural border in the chick by rostral endoderm signalling. Development 139, 1034–1044.

Sasai, N., Mizuseki, K., Sasai, Y., 2001. Requirement of FoxD3-class signaling for neural crest determination in Xenopus. Development 128, 2525–2536.

Sato, S., Ikeda, K., Shioi, G., Ochi, H., Ogino, H., Yajima, H., Kawakami, K., 2010. Conserved expression of mouse Six1 in the pre-placodal region (PPR) and identification of an enhancer for the rostral PPR. Dev Biol 344, 158–171.

Schlesinger, S., Meshorer, E., 2019. Open Chromatin, Epigenetic Plasticity, and Nuclear Organization in Pluripotency. Dev Cell 48, 135–150.

Senft, A.D., Bikoff, E.K., Robertson, E.J., Costello, I., 2019. Genetic dissection of Nodal and Bmp signalling requirements during primordial germ cell development in mouse. Nat Commun 10, 1089.

Shaffer, A.L., Lin, K.I., Kuo, T.C., Yu, X., Hurt, E.M., Rosenwald, A., Giltnane, J.M., Yang, L., Zhao, H., Calame, K., Staudt, L.M., 2002. Blimp-1 orchestrates plasma cell differentiation by extinguishing the mature B cell gene expression program. Immunity 17, 51–62.

Sheng, G., Stern, C.D., 1999. Gata2 and Gata3: novel markers for early embryonic polarity and for nonneural ectoderm in the chick embryo. Mech Dev 87, 213–216.

Simoes-Costa, M., Bronner, M.E., 2013. Insights into neural crest development and evolution from genomic analysis. Genome Res 23, 1069–1080.

Simoes-Costa, M., Bronner, M.E., 2015. Establishing neural crest identity: a gene regulatory recipe. Development 142, 242–257.

Simoes-Costa, M.S., McKeown, S.J., Tan-Cabugao, J., Sauka-Spengler, T., Bronner, M.E., 2012. Dynamic and differential regulation of stem cell factor FoxD3 in the neural crest is Encrypted in the genome. PLoS Genet 8, e1003142.

Stern, C.D., Ireland, G.W., 1981. An integrated experimental study of endoderm formation in avian embryos. Anat Embryol (Berl) 163, 245–263.

Streit, A., 2018. Specification of sensory placode progenitors: signals and transcription factor networks. Int J Dev Biol 62, 195–205.

Streit, A., Berliner, A.J., Papanayotou, C., Sirulnik, A., Stern, C.D., 2000. Initiation of neural induction by FGF signalling before gastrulation. Nature 406, 74–78.

Streit, A., Lee, K.J., Woo, I., Roberts, C., Jessell, T.M., Stern, C.D., 1998. Chordin regulates primitive streak development and the stability of induced neural cells, but is not sufficient for neural induction in the chick embryo. Development 125, 507–519.

Streit, A., Sockanathan, S., Perez, L., Rex, M., Scotting, P.J., Sharpe, P.T., Lovell-Badge, R., Stern, C.D., 1997. Preventing the loss of competence for neural induction: HGF/SF, L5 and Sox-2. Development 124, 1191–1202.

Streit, A., Stern, C.D., 1999. Establishment and maintenance of the border of the neural plate in the chick: involvement of FGF and BMP activity. Mech Dev 82, 51–66.

Streit, A., Stern, C.D., 2001. Combined whole-mount in situ hybridization and immunohistochemistry in avian embryos. Methods 23, 339–344.

Strobl-Mazzulla, P.H., Sauka-Spengler, T., Bronner-Fraser, M., 2010. Histone demethylase JmjD2A regulates neural crest specification. Dev Cell 19, 460–468.

Stuhlmiller, T.J., García-Castro, M.I., 2012. FGF/MAPK signaling is required in the gastrula epiblast for avian neural crest induction. Development 139, 289–300.

Surani, M.A., Hayashi, K., Hajkova, P., 2007. Genetic and epigenetic regulators of pluripotency. Cell 128, 747–762.

Teng, L., Mundell, N.A., Frist, A.Y., Wang, Q., Labosky, P.A., 2008. Requirement for Foxd3 in the maintenance of neural crest progenitors. Development 135, 1615–1624.

Theunissen, T.W., Jaenisch, R., 2017. Mechanisms of gene regulation in human embryos and pluripotent stem cells. Development 144, 4496–4509.

Trevers, K.E., Prajapati, R.S., Hintze, M., Stower, M.J., Strobl, A.C., Tambalo, M., Ranganathan, R., Moncaut, N., Khan, M.A.F., Stern, C.D., Streit, A., 2017. Neural induction by the node and placode induction by head mesoderm share an initial state resembling neural plate border and ES cells. Proc. Nat. Acad. Sci. USA 115, 355–360.

Uchikawa, M., Ishida, Y., Takemoto, T., Kamachi, Y., Kondoh, H., 2003. Functional analysis of chicken Sox2 enhancers highlights an array of diverse regulatory elements that are conserved in mammals. Dev Cell 4, 509–519.

Vincent, S.D., Dunn, N.R., Sciammas, R., Shapiro-Shalef, M., Davis, M.M., Calame, K., Bikoff, E.K., Robertson, E.J., 2005. The zinc finger transcriptional repressor Blimp1/Prdm1 is dispensable for early axis formation but is required for specification of primordial germ cells in the mouse. Development 132, 1315–1325.

Wamaitha, S.E., Niakan, K.K., 2018. Human Pre-gastrulation Development. Curr Top Dev Biol 128, 295–338.

Yu, J., Angelin-Duclos, C., Greenwood, J., Liao, J., Calame, K., 2000. Transcriptional repression by blimp-1 (PRDI-BF1) involves recruitment of histone deacetylase. Mol Cell Biol 20, 2592–2603.

