## Supplementary Figures for "PRDM1 controls the sequential activation of neural, neural crest and sensory progenitor determinants by regulating histone modification"

Supplemental Figures

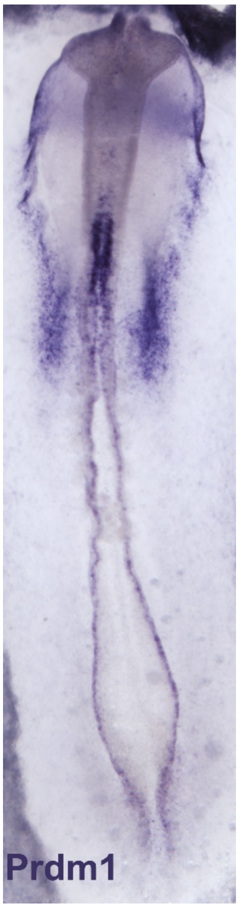


**Supplemental Figure 1. *PRDM1* expression at the 10-somite stage.** *PRDM1* expression is absent in sensory placodes, except for epibranchial precursors.

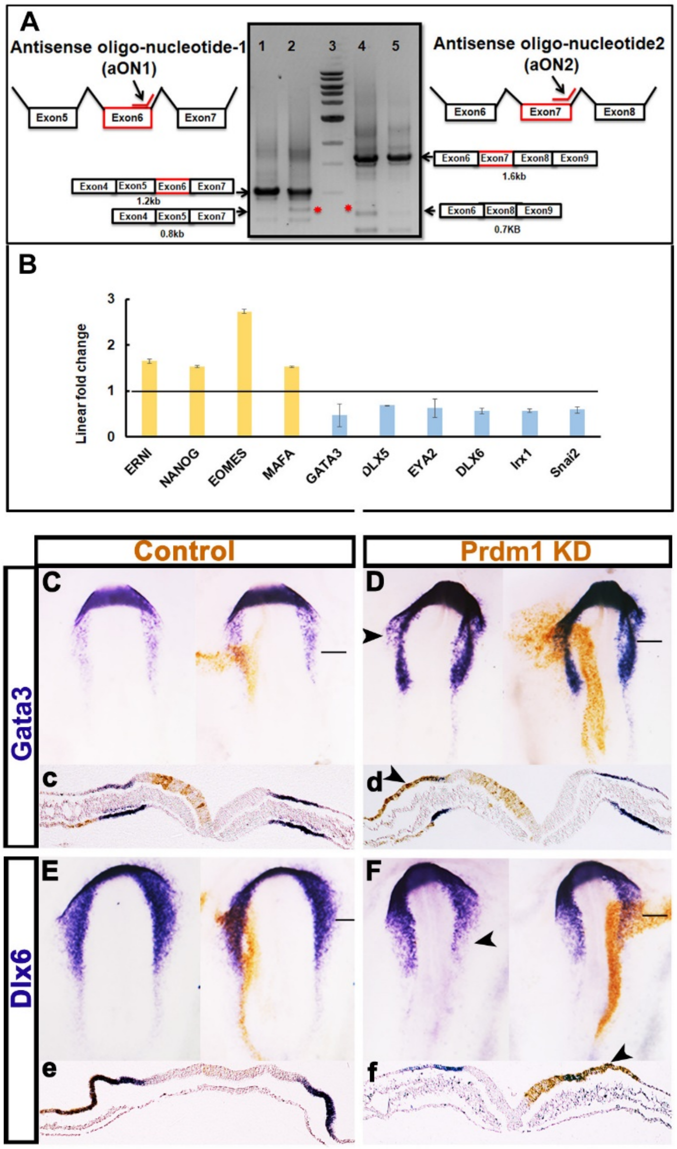


**Supplemental Figure 2.**

A. Two antisense oligonucleotides were designed to knockdown PRDM1. aON1 targets the boundary of exon 6 and intron 6. RT PCR of control (1) and aON1 targeted tissue shows both wildtype and exon 6 deletion (*). aON2 targets exon 7- intron 6 boundary leading to exon 7 deletion. RT PCR shows both wildtype and exon 7 deleted products.

B. Bar diagram showing NanoString nCounter data for selected genes. PRDM1 was knocked down using a combination of aON1 and aON2 at early primitive streak stages; sensory progenitors were harvested at headfold stages and analysed by NanoString. Pluripotency genes are upregulated, while neural plate border, sensory progenitor and some neural crest genes are downregulated as compared to controls.

C.- F. PRDM1 knockdown leads to loss of neural plate border markers *Gata3* and *Dlx6.* Control (C, c, E, e) or PRDM1 aON1/2 (D, d, F, f) were electroporated at early primitive streak stages (brown). Embryos were fixed at headfold or early somite stages and gene expression was assessed by in situ hybridisation. While the expression of *Gata3* and *Dlx6* is normal in controls, expression is lost in PRDM1 knockdowns (D, d, F,f; arrowheads). Panels on the left show embryos prior to visualisation of ON targeted cells by fluorescein immunolabelling (brown); c-f show sections through the same embryos shown in C-F at the level of the black line.

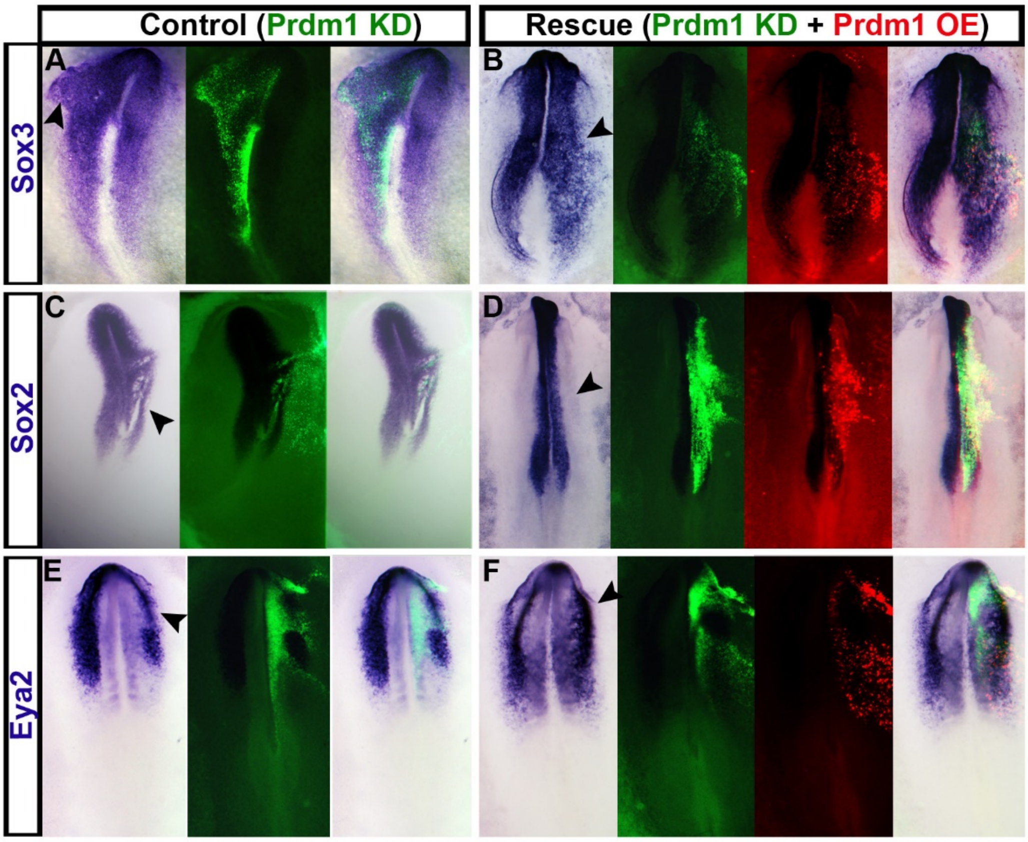


**Supplemental Figure 3. Rescue of PRDM1 knockdown.**
A, C, E. PRDM1 knockdown using aONs leads to expansion of *Sox3*, and downregulation of *Sox2* and *Eya2* (arrowheads). Images on the left show in situ hybridisation, middle panels cells targeted with fluorescein labelled aONs and images on the right show overlay. B, D, F. Coelectroporation of PRDM1 and PRDM1 aONs restores normal gene expression (arrowheads). Panels from left to right show: in situ hybridisation, fluorescein labelled aONs, PRDM1 expression construct (pCAB PRDM1-IRES-RFP) and overlay.

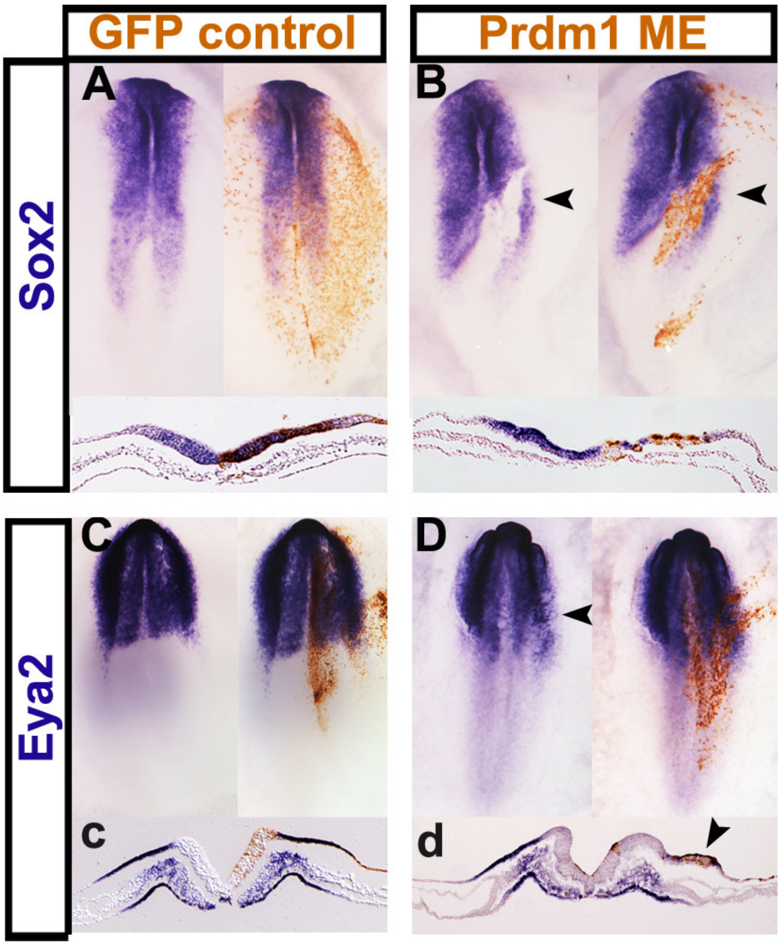


**Supplemental Figure 4. Misexpression of PRDM1 at late primitive streak stages lead to repression of neural and sensory progenitor markers.**
GFP controls (A, a, C, c) or PRDM1-IRES-RFP (B, b, D, d) were electroporated into primitive streak stage embryos. At headfold stages embryos were assess for *Sox2* and *Eya2* expression by in situ hybridisation. Gene expression is normal in controls, however, PRDM1 misexpression results in downregulation of *Sox2* and *Eya2* (arrowheads). a-d show sections through the same embryos as shown in A-D.
